# Evolution of form I-related RuBisCOs revisited: Uncertainties and diversity encompassing Bacteria and Archaea

**DOI:** 10.64898/2026.09.21.753380

**Authors:** Kohei Bamba, Euki Yazaki, Takuro Nakayama, Yuji Inagaki

## Abstract

Forms I′, I″, and Iα RuBisCOs were recently identified in bacterial metagenome-assembled genomes (MAGs) and are significant for evolutionarily bridging form I, which is responsible for the major part of carbon fixation on Earth, and other forms. Pioneering studies focused mainly on the biochemical and structural characteristics of “form I-related” RuBisCOs and deduced the evolution of this protein family over phylogenetic relationships among form I and form I-related sequences. We here surveyed form I-related sequences in public databases and conducted phylogenetic analyses incorporating the updated diversity of form I-related RuBisCOs. This study yielded two major outcomes regarding RuBisCO evolution. First, the analyses presented here suggest uncertainties overlooked in the previously published studies. It appeared to be difficult to conclude (i) the phylogenetic relationship among forms I, I′, and I″, (ii) the monophyly of form I′, or (iii) the monophyly of form Iα. Second, we identified a gene encoding form Iα RuBisCO in an archaeal MAG, and this protein is likely involved in the pentose bisphosphate pathway. The results presented here suggest that the evolutionary route leading to form I in RuBisCO evolution remains unsettled.

## Introduction

RuBisCO, ribulose-1,5-bisphosphate carboxylase/oxygenase, is the most abundant enzyme on Earth and plays a major role in carbon fixation (Erb and Zarzycki 2018; Bar-On and Milo 2019; Prywes et al. 2023). This protein yields 3-phosphoglycerate (3-PGA) from carbon dioxide and ribulose-1,5-bisphosphate (RuBP), a key step in the Calvin-Benson-Bassham (CBB) cycle (Prywes et al. 2023). The RuBisCO protein family comprises distinct forms whose structures, particularly quaternary structure, and distributions in the tree of life differ from one another (Tabita 1999; Tabita et al. 2008). Form I RuBisCO has been found in cyanobacteria and diverse photosynthetic eukaryotes that bear plastids traced to a cyanobacterial endosymbiont (Watson and Tabita 1997). The tertiary structure of form I RuBisCO is composed of eight large and eight small subunits—the large subunits form the catalytic core capped by the small subunits (Andersson and Taylor 2003). Form II RuBisCO has been documented in chemotrophic bacteria and in a restricted lineage of photosynthetic eukaryotes (i.e., dinoflagellates), which acquired form II genes horizontally from a bacterium (Watson and Tabita 1997). No small subunit is required for form II RuBisCO, and it exists as a dimer or a hexamer (Schneider et al. 1986; Satagopan et al. 2014; Arbing et al. 2016). In Archaea, neither form I nor II has been found, but form III RuBisCO, which lacks any small subunit and can be assembled by multiple (five at most) dimers of the protein (Maeda et al. 1999; Kitano et al. 2001; Huang and Szebenyi 2023). The vast majority of form III RuBisCO sequences have been identified in Archaea, but uncultivated bacteria belonging to the candidate phyla radiation (CPR) were also known to bear RuBisCOs related to this particular form (Wrighton et al. 2016). Although form III RuBisCO catalyzes the carboxylation of RuBP as other RuBisCOs do (Ezaki et al. 1999), this appeared to be involved in the pentose bisphosphate (PBP) pathway unique to Archaea, instead of the CBB pathway (Aono et al. 2015; Sato, Atomi, and Imanaka 2007; Aono et al. 2012). Besides the three forms of RuBisCO described above, there are RuBisCO-like proteins (RLP or form IV), which resemble the above-mentioned RuBisCO forms structurally but do not catalyze carbon fixation (Ashida et al. 2003; Hanson and Tabita 2003; Li et al. 2005).

Recent studies have expanded our understanding of the diversity of RuBisCO and narrowed gaps between form I and other forms. So far, three clades of RuBisCO sequences, which bear a specific evolutionary affinity to, but are clearly distinct from, the large subunit of form I sequences, have been identified. Henceforth, we designate the RuBisCO sequences mentioned above collectively as “form I-related” RuBisCOs. Form I′ RuBisCO was first identified from metagenome-assembled genomes (MAGs) derived from bacteria belonging to the phylum Chloroflexi (Banda et al. 2020). Then, two additional clades of form I-related RuBisCOs, forms I″ and Iα, were reported (West-Roberts et al. 2021; Liu et al. 2023). Unlike form I RuBisCO, form I′, I″, or Iα RuBisCO requires no small subunit. There are experimentally determined tertiary and quaternary structures of forms I′ and I″ RuBisCOs, and both appeared to be homo-octameric structures (Banda et al. 2020; Schulz et al. 2022; Liu et al. 2023). In contrast, despite the lack of direct investigation of the tertiary/quaternary structure, multiple studies consistently suggested a dimeric assembly of form Iα RuBisCO (Schulz et al. 2022; Liu et al. 2023). The previously published phylogenies of form I and form I-related RuBisCOs (rooted by other forms) consistently grouped forms I, I′, and I″ sequences together, excluding form Iα sequences (West-Roberts et al. 2021; Schulz et al. 2022; Liu et al. 2023). Nevertheless, the branching order among forms I, I′, and I″ RuBisCOs appeared to be incongruent across the studies. Schulz et al. (2022) discussed the evolution of form I-related RuBisCOs based on the tree in which form I was closer to form I″ than form I′. On the other hand, Liu et al. (2023) elaborated on interpreting the differences among forms I, I′, and I″ RuBisCOs based on the tree uniting forms I and I′, rather than form I″.

Form I-related RuBisCOs are indeed significant in inferring the evolutionary trajectory from ancestral forms (i.e., forms II and III) to form I. Nevertheless, we need to pay close attention to whether the evolutionary relationship underlying the scenario for RuBisCO evolution is valid and whether any alternatives can be ruled out—this aspect, in principle, is the matter of molecular phylogeny. It is well known that various aspects of phylogenetic analyses can (sometimes severely) bias inferences. One of the key aspects in phylogeny for better inferences is which sequences/taxa to include in an alignment, so-called “taxon sampling” (Heath, Medtke and Hillis 2008; Nabhan and Sarkar 2011). Even with the most realistic substitution model available and a rigorous method for tree reconstruction, a false tree can be inferred from an alignment in the absence of a particular key sequence/group of sequences (e.g., Yazaki et al. 2021).

On the contrary, the inclusion of a problematic sequence/group of sequences, e.g., extremely rapidly evolving sequences of which the tempo and mode of substitutions depart largely from those assumed in the substitution model specified (i.e., model misspecification), can result in a false inference (e.g., Isogai et al. 2025). It is also important to evaluate the robustness of the tree topology of interest. A particular phylogenetic relationship reconstructed from an alignment does not necessarily mean we can ignore an alternative relationship that was not reconstructed. Non-parametric bootstrap and other methods have been widely used to assess the reliability of bipartitions in the tree inferred from the alignment of interest (Felsenstein 1985; Guindon et al. 2010; Minh et al. 2013; Hong et al. 2018). Nevertheless, in a strict sense, we cannot judge whether alternative evolutionary routes are possible based on the above-mentioned support values for bipartitions. In the case of an alternative tree topology (or topologies) competing with the tree inferred from a phylogenetic alignment, an approximately unbiased test or similar tests can directly examine whether the alternative one can be discarded at a certain statistical level (Kishino and Hasegawa 1989; Shimodaira and Hasegawa 1999; Shimodaira 2002). With the methodological points in phylogenetics described above in mind, we here expanded the sequence diversity of form I-related RuBisCOs and rigorously examined the branching order of form I-related RuBisCOs.

In this study, we first identified novel forms I′ and Iα RuBisCO sequences in MAGs. The newly identified RuBisCO sequences, together with those analyzed in Liu et al. (2023), were subjected to phylogenetic analyses. We prepared and analyzed a new RuBisCO alignment considering a broader range of form I-related sequences than those analyzed in the pioneering studies. The analyses presented below unveiled potential uncertainties in the evolution of form I and form I-related RuBisCOs that have not previously been questioned, e.g., the monophyly of form Iα sequences. In addition, the RuBisCO phylogeny appeared to be insufficient to resolve the relationship among forms I, I′, and I″ with confidence, blurring when the C-terminal extension found in forms I and I″ (but not in form I′) emerged. Finally, we report the first archaeal form Iα RuBisCO, which is likely involved in the PBP pathway.

## Materials & Methods

### Retrieval of RuBisCO sequences from public databases

We retrieved 333 RuBisCO amino acid sequences analyzed in Erb et al. (2012) from the National Center for Biotechnology Information by using their GI numbers. 31 out of the 333 RuBisCO sequences analyzed in Erb et al. (2012) were manually selected and used as queries during the homology search described below. We searched for novel RuBisCO sequences in the NCBI nr, refseq, and env_nr databases (downloaded on June 16, 2022) using the blastp program in BLAST+ v.2.13.0. The homology search was restricted to the archaeal and bacterial sequences. Based on the reports from the BLAST searches, we retrieved 13,195 RuBisCO sequences that satisfied both criteria described below, namely (i) *E*-value equal to or less than 1×10^-10^ and (ii) query coverage per HSP equal to or greater than 50.

### Phylogenetic Analyses

We prepared and analyzed two RuBisCO alignments, “all_forms_1736” and “fI_related_266.” Both alignments comprise subsets of the RuBisCO sequences identified in this study (see above) and those analyzed in Liu et al. (2023). The analysis of all_forms_1736 aimed to identify previously uncharacterized RuBisCO sequences related to form I. We then analyzed fI_related_266 to closely assess the evolution of form I and form I-related RuBisCOs. The detailed procedures for preparing the two RuBisCO alignments and subsequent phylogenetic analyses of them are described in the following sections.

#### Analysis of all_forms_1736

We reduced the sequence redundancy in the 13,195 RuBisCO sequences retrieved by the initial blast search (see above) by using CD-HIT v.4.8.1 (Huang et al. 2010) with “-c 0.7 -n 5” option, and the resultant 1,501 RuBisCO sequences were aligned with the 235 sequences analyzed in Liu et al. (2023) by MAFFT v.7.505 (Katoh and Standley 2013) under the L-INS-i model. The initial alignment of the 1,736 RuBisCO sequences was trimmed using BMGE v.2.0 (Criscuolo and Gribaldo 2010) with default settings, retaining 239 amino acid positions for subsequent phylogenetic analyses. The maximum likelihood (ML) phylogenetic analysis of all_forms_1736 was conducted by IQ-TREE2 v.2.2.0 (Minh et al. 2020) under the LG + F + R10 model selected by ModelFinder (Kalyaanamoorthy et al. 2017). Due to the computational burden associated with the alignment size, we did not apply any site-heterogeneous model to the ML analysis. Support values for bipartitions in the ML tree were calculated by UFBoot2 (Hoang et al. 2018) implemented in IQ-TREE with 1,000 replicates (ultrafast bootstrap support values; UFBPs).

#### Analyses of fI_related_266

The ML analysis of all_forms_1736 identified 31 sequences regarded as form I-related RuBisCOs that were not analyzed in Liu et al. (2023). As done to prepare all_forms_1736, the newly identified form I-related sequences and those analyzed in Liu et al. (2023) were put together and aligned by MAFFT under the L-INS-i model. The aligned 266 RuBisCO sequences were subjected to BMGE with default settings to select 394 positions for phylogenetic analyses, yielding the final form I-related RuBisCO alignment (fI_related_266). First, fI_related_266 was subjected RAxML-ng v2.0.2 with the option “pars{100}, rand{100},” in which tree search started from 100 parsimony trees and 100 randomly generated trees, under the LG + Γ + F model. For this first analysis, we assigned the “site homogeneous” LG + Γ + F model, which disregarded heterogeneity in amino acid composition across alignment positions. Instead, a tree search was initiated from 200 starting trees to cover a broad tree space. The tree yielded from the RAxML analysis was then supplied to the second analysis using IQ-TREE v 3.1.3 (Minh et al. 2020) as the starting tree. In the IQ-TREE analysis, the substitution model was set to a more realistic LG + C60 + F + R7 model, which accounts for heterogeneity in amino acid composition across alignment positions. The substitution model was selected by ModelFinder with “-mset LG+C60” option. To calculate support values for bipartitions in the ML tree, we conducted UFboot2 (1,000 replicates), the non-parametric bootstrap method (100 replicates), and AUTOEB v.1.0 with default settings (Banba, Harada and Inagaki 2024). In the non-parametric bootstrap analysis, the parameters for compositional heterogeneity of amino acids among sites in the alignment were calculated by the PMSF model (Wang et al. 2018) over the ML tree.

### Protein Structure Prediction

We predicted the quaternary structures of two form Iα RuBisCOs of a *Candidatus* Bathyarchaeota archaeon (GenBank protein accession number, MCJ7457314) and the deltaproteobacterium *Dissulfurimicrobium hydrothermale* (UKL14116.1) by AlphaFold2 (Yang et al. 2023). For each prediction, we submitted a FASTA file containing eight identical amino acid sequences of interest to AlphaFold2 in multimer mode. Separately, we ran AlphaFold2 to predict the quaternary structure from eight amino acid sequences of form I′ RuBisCO of *Candidatus* Prominefilum breve, of which the RuBisCO octamer was determined experimentally (PDB accession number 6URA).

## Resutls

To re-evaluate the diversity of form I-related RuBisCOs, we retrieved RuBisCO sequences from the GenBank database. According to the phylogenetic analysis of the alignment containing the retrieved sequences (i.e., all_forms_1736; see above), the vast majority of the retrieved RuBisCO sequences were considered form II or III, as they did not form a clade with the previously known form I, I′, I″, or Iα. Briefly, none of the RuBisCO sequences identified in this study fell into the clade of form I or I″, but 27 and four branched with the previously known form I′ and Iα sequences, respectively (see below). We prepared and phylogenetically analyzed a RuBisCO alignment designated as “fI_related_266,” comprising the newly identified 31 sequences and those analyzed in Liu et al. (2023).

In the ML tree inferred from fI_related_266, all of the form I and form I-related RuBisCO sequences were grouped with a UFBP of 100% and an MLBP of 100%, excluding the RuBisCO sequences belonging to other forms (Fig. 1A). The monophyly of form I sequences and that of form I″ sequences were recovered with full statistical support (schematically shown in orange and green triangles; Fig. 1A). AUTOEB examined the node uniting 80 form I sequences and that uniting three form I″ sequences. This program runs an approximately unbiased (AU) test (Shimodaira and Hasegawa 1999; Shimodaira 2002) comparing the original (ML) and the two alternative trees generated from the ML tree by nearest-neighbor interchange (NNI) on the bipartition of interest (designated as NNI-alternative trees). For the two nodes examined here, the AU test rejected both NNI-alternative trees, and we thus regarded them as “resolved.”

**Figure 1.**
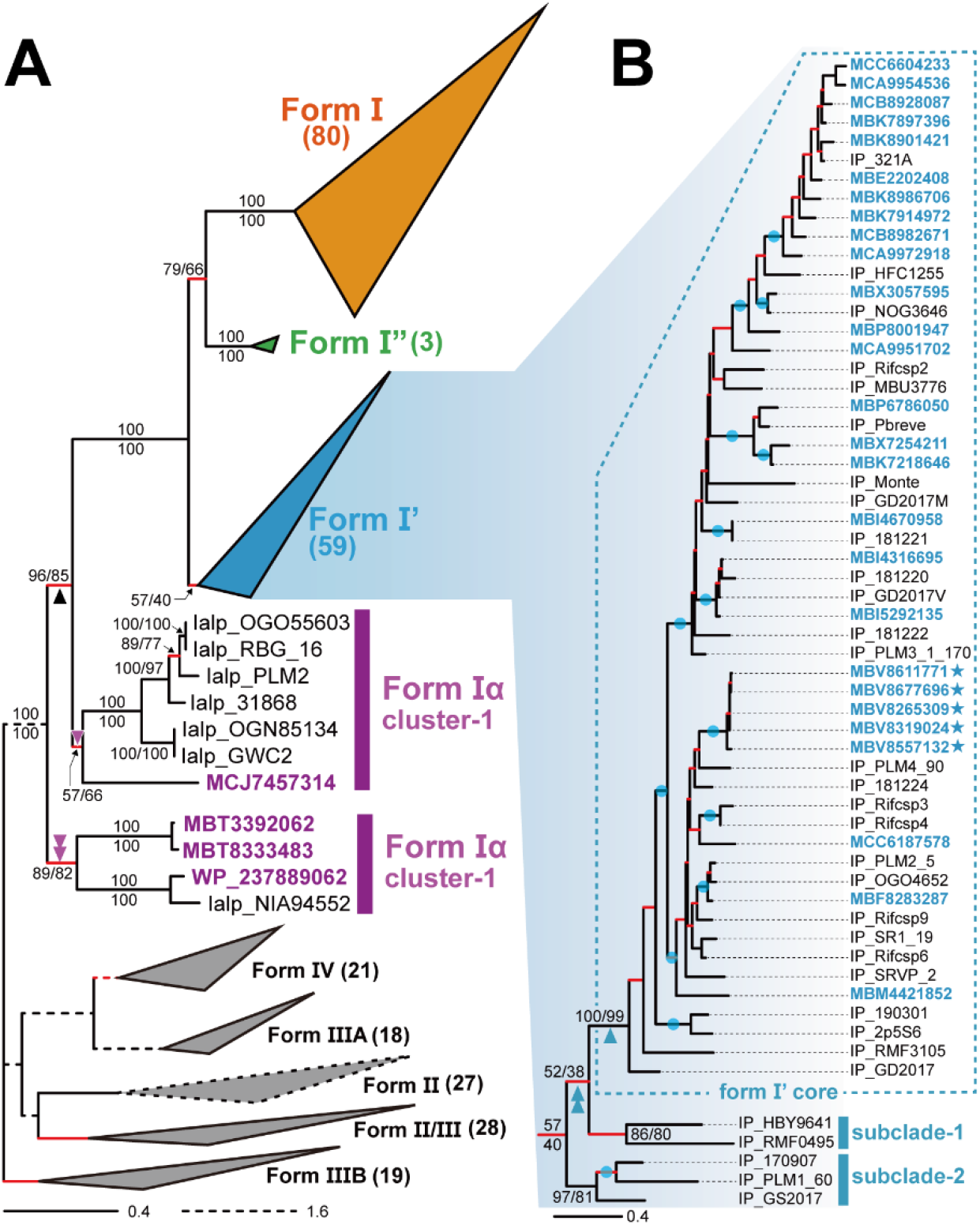
RuBisCO phylogeny. **(A)** The overall phylogenetic relationship among 80 form I, 59 form I′, three form I″, 11 form 1α, and other forms was inferred by the ML method. Major clades are schematically shown as triangles (the number of sequences composed of each clade is shown in parentheses). Form Iα sequences identified in this study are colored in purple. Support values from UFBoot2 (UFBPs) and those from the ML bootstrap analysis (MLBPs) are presented above and below the corresponding bipartitions, respectively, or on the left and right of slashes, respectively. The bipartitions colored in red were regarded as “unresolved” by AUTOEB. **(B)** The detail of the form I′ clade. Form I′ sequences identified in this study are colored in blue. The sequences identified in the MAGs of Planctomycota are marked by stars. UFBPs and MLBPs for the bipartitions important for assessing the monophyly of form I′ RuBisCO are shown in the same way as in (A). Other than those, only bipartitions that received both UFBP and MLBP ≥90% are marked by dots. The nodes colored in red were regarded as “unresolved” by AUTOEB. This figure was created using TreeMage (https://github.com/Funny-Silkie/TreeMage)

The clade of the 32 previously reported form I′ and 27 newly identified form I-related sequences was recovered (schematically shown in a blue triangle; Fig. 1A), albeit the support was rather weak, namely a UFBP of 57% and an MLBP of 40%. The node uniting form I′ and the 27 newly identified form I-related sequences was regarded as “unresolved” by AUTOEB (marked in red; Fig. 1A). Four of the newly identified form I-related sequences were excluded from the cluster of Forms I, I″, and I′ and branched with the previously reported form Iα sequences (Fig. 1A).

In the following sections, we evaluate (i) the monophyly of form I′ sequences, (ii) the relationship among forms I, I′, and I″, and (iii) the monophyly of form Iα sequences, in light of the RuBisCO phylogeny incorporating the 31 form I-related sequences identified in this study.

### Monophyly of form I′ RuBisCO (or not?)

The 27 form I-related RuBisCO sequences identified in this study fell within the diversity of the form clade I′ sequences reported in a previous study (Fig. 1B). In particular, the node uniting the 27 form I′ sequences reported in a previous study (Liu et al. 2023) and the 27 form I-related sequences identified in this study received a UFBP of 100% and an MLBP of 99% (highlighted by a blue arrowhead in Fig.1B). The node mentioned above was judged as “resolved” by AUTOEB. We conclude here that the 27 newly identified form I-related sequences belong to form I′. We here designate the subclade comprising the 54 form I′ sequences discussed above as “form I′ core.” All of the previously reported form I′ RuBisCOs were from bacteria belonging to Chloroflexi (Liu et al. 2023). Likewise, 22 out of the 27 newly identified form I′ sequences were of bacteria belonging to Chloroflexi, except for five of Planctomycetota (marked by stars in Fig. 1B). Form I′ core appeared to be appended by “subclade-1” and “subclade-2” in this order (Fig. 1B): Two previously reported sequences, IP_RMF0495 and IP_HBY9641, were united with a UFBP of 86% and an MLBP of 80%, forming subclade-1. The remaining three sequences, IP_PLM1_60, IP_170907, and IP_GS2017, formed subclade-2 with a UFBP of 97% and an MLBP of 81%.

The monophyly of form I′ RuBisCO was recovered as the ML estimate, but a potential paraphyly of this RuBisCO form appeared to be difficult to exclude. First, based on the statistical support, the monophyly of the 59 form I′ sequences is still arguable (see above; Figs. 1A & B). In addition, the relationship among form I′ core, subclade-1, subclade-2, and other RuBisCO sequences remained inconclusive. The grouping of form I′ core and subclade-1 received only weak statistical support, a UFBP of 52% and an MLBP of 38%. Likewise, AUTOEB comparing the ML tree and two NNI-alternative trees, which were generated by NNI on the node marked by a blue double-arrowhead, regarded as “unresolved,” as neither of the two NNI-alternative trees was rejected at the 5% level of statistical confidence (marked in red; Fig. 1B).

To examine the evolution of form I′ sequences more comprehensively, we conducted an extra AU test assessing 15 test trees representing all possible relationships among (i) form I′ core, (ii) form I′ subclade-1, (iii) form I′ subclade-2, (iv) the clade of forms I and I″, and (v) other RuBisCO sequences. Among the 15 test trees, there were three trees with the monophyly of form I′ (Trees 1-3; Tree 1 corresponds to the ML tree; see the left panel of Fig. 2). Significantly, only five out of Trees 4-15, in which form I′ was paraphyletic, were rejected at the 5% level of statistical confidence (the right panel of Fig. 2). Thus, it is too early to assume that the currently identified form I′ sequences were derived from a single ancestral RuBisCO sequence. To judge whether form I′ is truly monophyletic, we need further surveys of form I-related RuBisCO sequences in natural environments and to conduct the phylogenetic analysis based on an alignment including a broader collection of form I′ sequences than that covered in the alignment assessed in this study.

**Figure 2.**
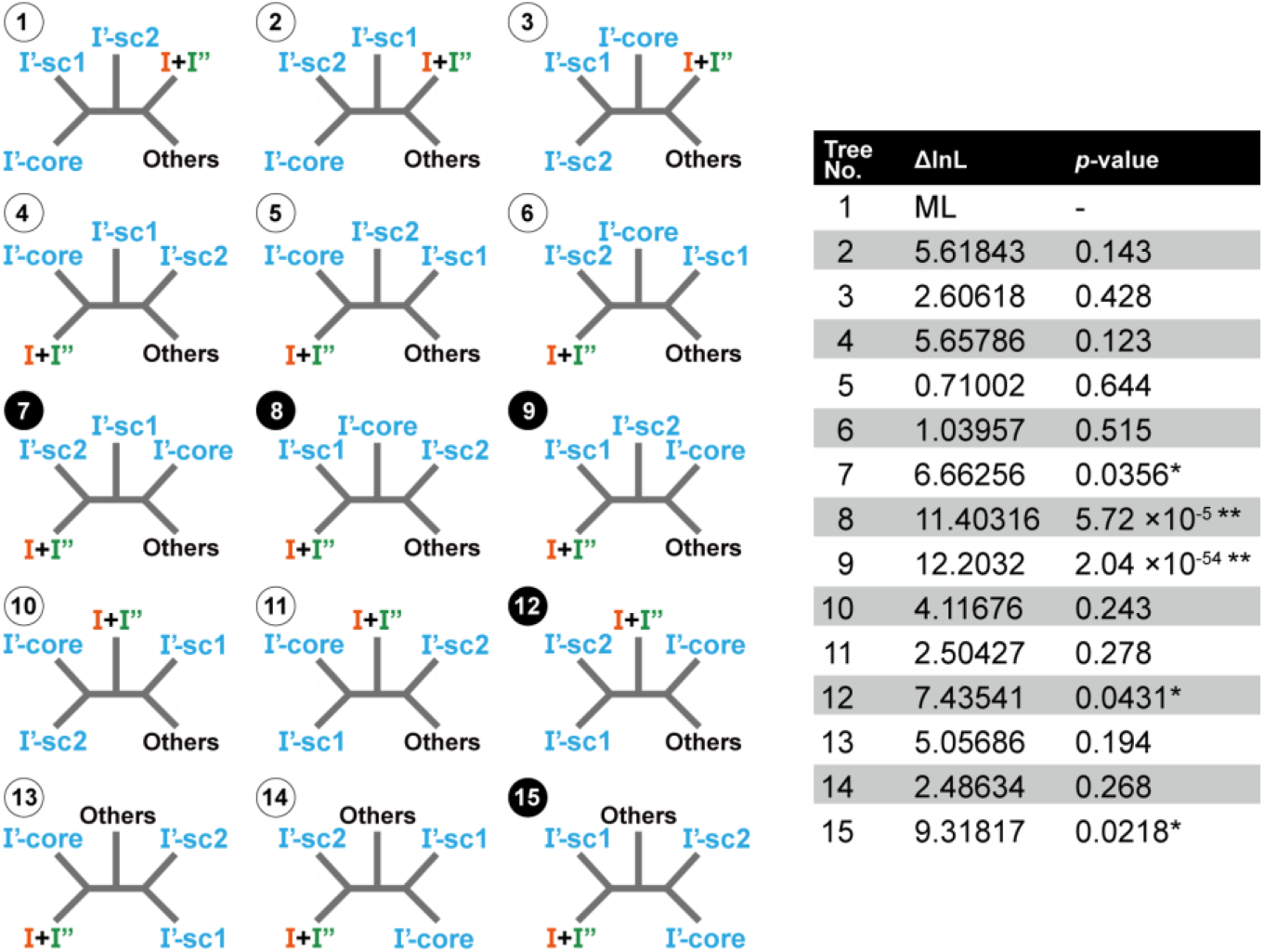
An AU test assessing the evolution of form I′ sequences. The 15 test trees subjected to the AU test are schematically shown in the left panel of the figure. They represent all possible relationships among (i) form I′ core (abbreviated as I′-core), (ii) form I′ subclade-1 (I′-sc1), (iii) form I′ subclade-2 (I′-sc2), (iv) the clade of forms I and I″ (I+I″), and (v) other RuBisCO sequences (Others). The table on the right summarizes the differences in log-likelihood (ΔlnL) between the ML tree (i.e., Tree 1) and each of Trees 2-15 and the *p*-values from the AU test. Trees 7, 12, and 15 were rejected at the 5% level of statistical confidence, marked with asterisks. Trees 8 and 9 were rejected at the 1% level of statistical confidence, marked with double asterisks.

### Uncertainty in the relationship among forms I, I′, and I″ RuBisCOs

As recovered in the RuBisCO phylogeny presented in Liu et al. (2023), forms I, I′, and I″ grouped together in the ML analysis with full statistical support and their monophyly was regarded as “resolved” by AUTOEB (Fig. 1A). On the other hand, the relationship among the three forms appeared to be different between Liu et al. (2023) and this study: Form I clade was joined with form I′ in the former but form I″ in the latter. The incongruence in the relationship among forms I, I′, and I″ between the two studies most likely stemmed from the lack of resolution in the RuBisCO alignment. In Liu et al. (2023), the sister relationship between forms I and I′ was supported by an MLBP of 73%. According to support values from an ML bootstrap analysis, there may be room for alternative branching patterns among the three forms, but this issue was not explicitly examined. We here recovered the grouping of forms I and I″ with a UFBP of 79% and an MLBP of 66% (Fig. 1A). The AU test assessing the relationship among forms I, I′ and I″ failed to reject the alternative tree corresponding to the ML tree recovered in Liu et al. (2023) in which form I and I′ were directly tied, excluding form I″ (NNI-1, *p* = 0.364; Fig. 3A). On the other hand, we regard that the direct union of forms I′ and I″ was much less likely, as “NNI-2” tree was barley rejected at the 5% level (*p* = 0.0495; Fig. 3A). In sum, it is currently difficult to conclude definitely whether form I′ or form I″ is the closest relative of form I.

**Figure 3.**
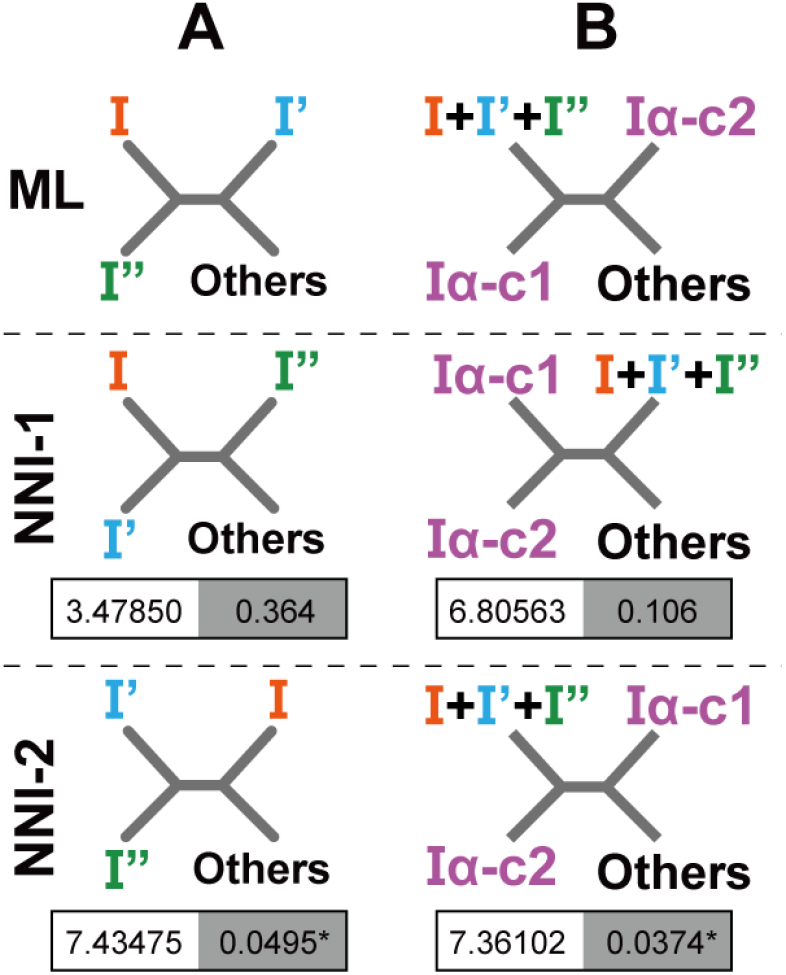
AU tests assessing the evolution of form I-related RuBisCOs. **(A)** Examinations for alternative relationships among forms I, I′, and I″. The ML tree, in which form I″ is closer to form I than form I′ (top), was compared to two alternative trees: “NNI-1,” in which form I is united directly to form I′ (center), and “NNI-2,” in which form I′ is united directly to form I″. Abbreviations: I, form I clade; I′, form I′ clade; I″, form I″ clade. For each alternative tree, the difference in log-likelihood (ΔlnL) from the ML tree and p-value are provided on the left and right in the box below the corresponding tree. NNI-2 alternative tree was rejected at the 5% level of statistical confidence (highlighted by an asterisk). The log-likelihood of the ML tree is -66167.19794. **(B)** Examinations for alternative relationships among the clade comprising forms I, I′, and I″, form Iα clan-1, and form 1α clan-2. The ML tree, in which the two clans of form Iα are paraphyletic (top), was compared to two alternative trees: “NNI-1,” in which the two clans of form Iα are monophyletic (center), and “NNI-2,” in which the two clans of form Iα are paraphyletic, and clan-2 branched directly with the clade of comprising forms I, I′, and I″ (bottom). Abbreviations: I+ I′+I″, the clade comprising forms I, I′, and I″; Iα-c1, form Iα clan-1; Iα-c2, form Iα clan-2. NNI-2 alternative tree was rejected at the 5% level of statistical confidence (highlighted by an asterisk). For each alternative tree, the difference in log-likelihood (ΔlnL) from the ML tree and p-value are shown as described in (A). The log-likelihood of the ML tree is -66170.50695.

### Revised diversity and evolution of form Iα RuBisCO

Forms I, I′, and I″ sequences excluded the seven form Iα sequences reported in Liu et al. (2023) and four form I-related RuBisCO sequences newly identified in this study (Fig. 1A). We here regard the four form I-related RuBisCO sequences identified in this study, which were related to but excluded from the clade of forms I, I′, and I″, as form Iα. In the ML phylogenetic analyses, the previously and newly identified form Iα sequences were coalesced into two clusters, clan-1 and clan-2 (Fig. 1A). In form Iα clan-1, six previously identified form Iα sequences grouped with full statistical support, and were then tied with a single newly identified form Iα sequence MCJ7457314 with a UFBP of 57% and an MLBP of 66% (Fig. 1A). Reflecting weak support values from UFBoot2 and ML bootstrap analysis, the particular node was regarded as “unresolved” by AUTOEB (see the bipartitions highlighted by a purple arrowhead; Fig. 1A). Form Iα clan-2 contained a single previously reported sequence (Ialp_NIA_0) and three newly identified ones (Fig. 1A). Within clan-2, Ialp_NIA_0 and UKL14116, and MBC8333483 and MBT3392062 tied with full statistical support (Fig. 1A). Clan-2 as a whole received somewhat high support values (i.e., a UFBP of 89% and an MLBP of 82%), but was regarded as “unresolved” by AUTOEB (see the bipartitions highlighted by a purple double-arrowhead; Fig. 1A).

If we accept the ML tree topology shown in Fig. 1A at the face value, form Iα RuBisCO is evolutionarily paraphyletic: Form Iα clan-1 and the clade of forms I, I′, and I″ sequences were connected directly to each other, excluding four of form Iα sequences composed clan-2 (Fig. 1A). Forms I, I′, I″ sequences, and seven out of the 11 form Iα sequences were grouped with a UFBP of 96% and an MLBP of 85% (marked by a black arrowhead in Fig. 1A). Notably, the possibility of the monophyly of form Iα remains. AUTOEB regarded the node separating the two clans paraphyletically as “unresolved” (highlighted by a black arrowhead; Fig. 1A), as the AU test failed to reject one of the two alternative trees, which represented the sister relationship between the two clans (NNI-1, *p* = 0.106; Fig. 3B). The third test tree, which was generated by substituting the positions of clan-1 and clan-2 in the ML tree, was rejected at the 5% statistical level (*p* = 0.0374; Fig. 3B). Thus, we need to be cautious about whether the phylogenetic relationship among the 11 form Iα sequences inferred reflects the truth. So far, based on the phylogenetic analyses conducted here, we cannot confidently deduce the ancestral position(s) of the 11 form Iα sequences relative to forms I, I′, and I″.

Six out of the seven previously identified form Iα sequences were commonly found in the MAGs derived from Chloroflexi bacteria. In this study, we identified additional form Iα sequences originating from the MAGs of Chloroflexi bacteria (GenBank protein accession nos. MBT3392062 and MBC8333483). In clan-2, Ialp_NIA_0, identified previously in a proteobacterial MAG (Liu et al. 2023), was revealed to be closely related to one of the newly identified sequences, UKL14116, found in the complete genome of the deltaproteobacterium *Dissulfurimicrobium hydrothermale* strain Sh68. The close phylogenetic affinity between two particular form Iα sequences implies that the “proteobacterium” harboring the form Iα sequence, Ialp_NIA_0, belongs to Deltaproteobacteria. Most notably, one of the new form Iα sequences (GenBank protein accession no. MCJ7457314) was identified in the MAG of a *Candidatus* Bathyarchaeota archaeon (GenBank accession number JALHTB010000210.1). We mapped the raw reads to the archaeal contig containing the form Iα gene to check whether its read depth departs from that of the entire contig. The mean depth of the form Iα gene was 48.9658 (minimum and maximum depths were 33 and 67, respectively), which is not substantially different from that of the entire contig (48.9610; minimum and maximum depths were 21 and 84, respectively). Thus, we found no strong evidence that the form Iα gene was misassembled into an archaeal genome fragment, although we cannot completely rule out the possibility that this RuBisCO sequence was artifactually incorporated into an archaeal MAG. If the entire contig is truly part of an archaeal genome, this form Iα sequence is the *de facto* first archaeal form I-related sequence known to date, demonstrating that the diversity of form I-related RuBisCOs encompasses both Bacteria and Archaea.

From the tertiary structural point of view, form Iα RuBisCO is distinct from form I′ or I″. Form Iα RuBisCO was predicted to work as a dimer, while both forms I′ and I″ are assembled into an octamer (Banda et al. 2020; Liu et al. 2023). We predicted the tertiary structures of two form Iα RuBisCOs identified in this study, one from *Candidatus* Bathyarchaeota archaeon and the other from the deltaproteobacterium *D. hydrothermale*, using AlphaFold2. Both predictions yielded dimeric structures of form Iα proteins (Fig. 4). Nevertheless, no close interaction between the dimers was suggested for either bacterial or archaeal form Iα protein examined (Fig. 4). In contrast, AlphaFold2 recovered the octameric structure of a bacterial form I′ RuBisCO (Fig. 4). The predictions conducted here further support the putative dimeric structure of form Iα RuBisCO proposed in Liu et al. (2023).

**Figure 4.**
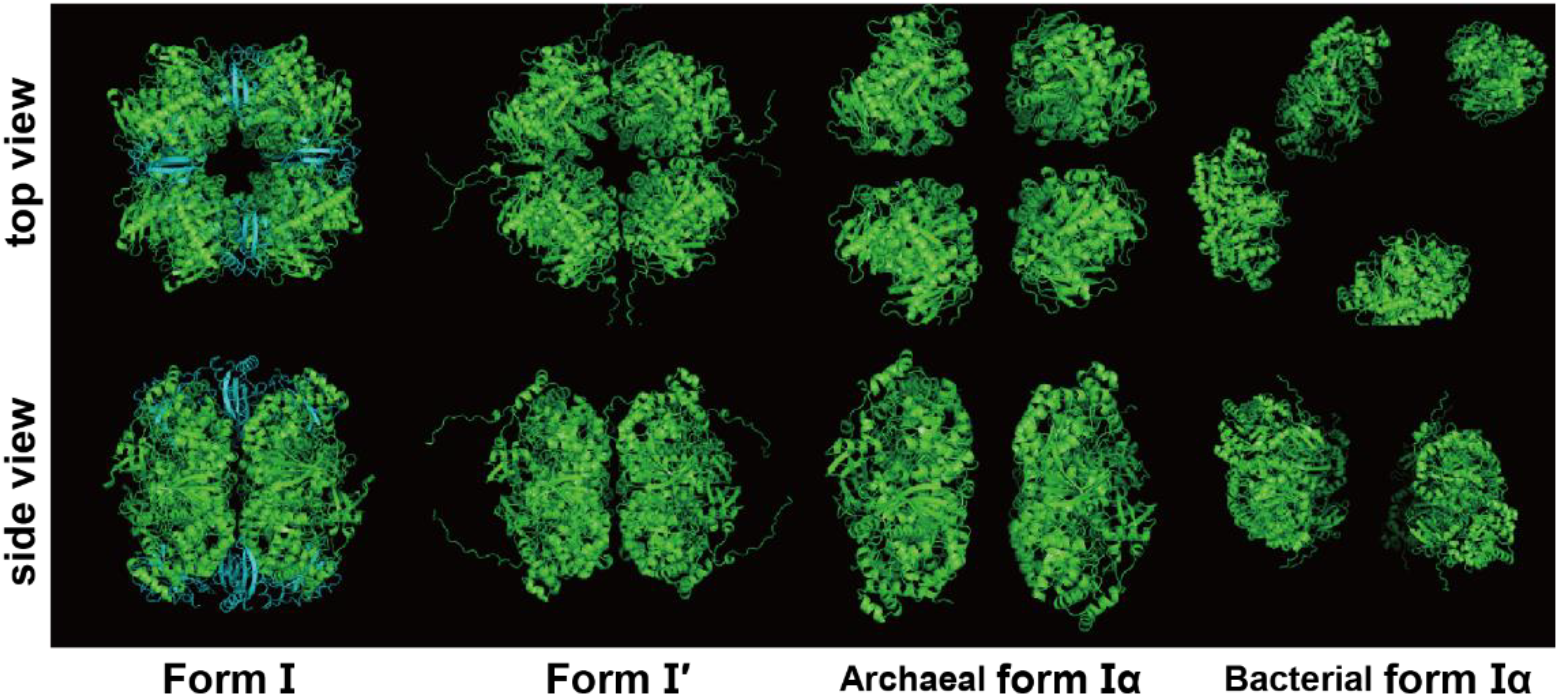
Quaternary structures of form I and form I-related RuBisCOs. Leftmost, the quaternary structure of eight large and eight small subunits of form I RuBisCO. The small- and large-subunit proteins are colored cyan and green, respectively. This structure was experimentally obtained by X-ray diffraction on RuBisCO proteins of the cyanobacterium *Synechococcus* sp. PCC6301 and deposited in the NCBI PDB under the accession number 1RBL. Note that the rest of the quaternary structures shown in this figure were predicted by AlphaFold2. Second, the predicted quaternary structure of eight peptides of form l′ RuBisCO of *Candidatus* Prominefilum breve. Third, the predicted quaternary structure of eight peptides of form lα RuBisCO of an archaeon belonging to *Candidatus* Bathyarchaeota. Rightmost, the predicted quaternary structure of eight peptides of form lα RuBisCO of the deltaproteobacterium *D. hydrothermale*. The quaternary structures were visualized using PyMOL (https://pymol.org/).

## Discussion

### An alternative scenario for the evolution of the C-terminal extensions found in forms I and I″ RuBisCOs

Liu et al. (2023) proposed a scenario for the C-terminal extension shared between form I and I″ RuBisCOs based on the phylogenetic relationship among forms I, I′, and I″. As the RuBisCO tree presented in Liu et al. (2023) united forms I and I′—the former bears the C-terminal extension, but the latter does not, while form I″ sequences have extensions at the C-termini. Thus, Liu et al. (2023) assumed that the ancestral molecule of forms I, I′, and I″ had already had the C-terminal extension, and form I′ lost it secondarily (Fig. 5A). Nevertheless, due to the uncertainty in the evolutionary relationship across forms I, I′, and I″, an alternative scenario should be considered. Our RuBisCO phylogeny (Fig. 1A) connected the two forms of RuBisCO bearing the C-terminal extension (i.e., forms I and I″) directly. This tree topology prompts us to propose an alternative scenario for the evolution of the C-terminal extension in forms I and I″ RuBisCO: A RuBisCO lacking the C-terminal extension is ancestral to the three forms, and after separating form I′ and the ancestral RuBisCO of forms I and I″, the latter acquired the C-terminal extension (Fig. 5B). Importantly, due to the uncertainty in the relationship among the three forms, we need to be cautious not to make any definite conclusion on the evolutionary trajectory of the C-terminal extensions of form I and I″ RuBisCOs.

**Figure 5.**
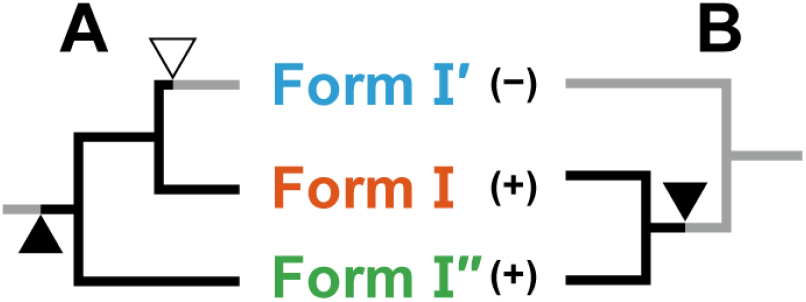
Scenarios for the evolution of the extensions at the C-termini of form I and I″ RuBisCOs. **(A)** The scenario proposed in Liu et al. (2023). This scenario was based on the RuBisCO tree assuming that form I′ is closer to form I than form I″. Thus, to explain the presence/absence of C-terminal extensions of the RuBisCO forms of interest (indicated by +/− in parentheses), it requires two events: (i) the acquisition of a C-terminal extension in the ancestral protein (indicated by a closed arrowhead), and (ii) the secondary loss of the extension of form I′ (indicated by an open arrowhead). **(B)** An alternative scenario under the RuBisCO tree in which form I″ is closer to form I than form I′. In the second scenario, the ancestral protein primarily lacked a C-terminal extension. Modern form I′ RuBisCO inherited this ancestral trait. The acquisition of a C-terminal tail occurred on the branch leading to the node uniting forms I and I″ (indicated by a closed arrowhead).

### Putative function of the archaeal form Iα RuBisCO

We investigated genes neighboring the form Iα RuBisCO gene in the archaeal MAG to obtain hints for which metabolic pathway this RuBisCO is involved in. In cyanobacterial genomes, the genes encoding the large and small subunits of form I RuBisCO, along with those encoding proteins involved in the CBB pathway, are found in close proximity, forming an operon. Likewise, the genes encoding form I-related RuBisCOs were found within the CBB operon in the corresponding genomes (Banda et al. 2020; Liu et al. 2023). As anticipated, all the bacterial form Iα RuBisCO genes newly identified appeared to be flanked by genes involved in the CBB pathway (Figs. 6A-C). In particular, the gene encoding phosphoribulokinase (PRK), which is involved exclusively in the CBB cycle, was found adjacent to the form Iα gene in each of the three bacterial MAGs identified in this study. In contrast, no gene involved in the CBB pathway was identified in the archaeal MAG that harbors the form Iα gene. To our surprise, tandem copies of the gene encoding ribulose-1,5-bisphosphate (R15P) isomerase, which comprises the pentose bisphosphate (PBP) pathway with form III RuBisCO in Archaea, were identified in one gene upstream of the form Iα gene in the MAG derived from *Candidatus* Bathyarchaeota archaeon (Fig. 6D). The PBP pathway has been found exclusively in Archaea, where R15P is isomerized to ribulose 1,5-bisphosphate (RuBP) by R15P isomerase, and RuBP is then carboxylated to 3-phosphoglycerate by form III RuBisCO. Thus, it is attractive to propose that the particular archaeon utilizes form Iα RuBisCO, instead of form III RuBisCO, to generate 3-phosphoglycerates through the carboxylation of RuBP.

**Figure 6.**
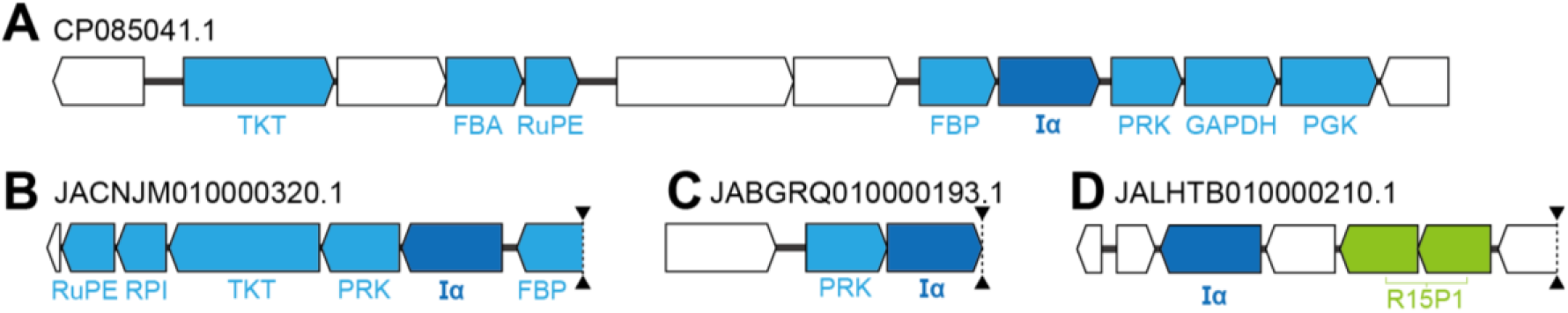
Protein genes flanking form Iα RuBisCO genes identified in this study. For each MAG/genome harboring a gene encoding form Iα RuBisCO, the protein genes surrounding a form Iα gene are schematically shown. The GenBank accession numbers for the genome/MAGs containing form Iα genes are provided individually. Form Iα genes (abbreviated as Iα) are colored in dark blue. Genes encoding enzymes involved in the CBB pathway are colored in light blue. **(A)** The form Iα gene in the complete genome of the deltaproteobacterium *Dissulfurimicrobium hy*drothermale strain Sh68 chromosome. The protein ID is UKL14116.1. **(B)** The form Iα gene in the MAG of an Anaerolineae bacterium NIOZ-UU35 NIOZ-UU_NODE_13407_length_7051_cov_12.9876. The protein ID is MBC8333483.1. **(C)** The form Iα gene in the MAG of a Chloroflexota bacterium isolate co234_bin83 NODE_32117_length_4173_cov_7.240408. The protein ID is MBT3392062.1. **(D)** The form Iα gene in the MAG of a *Candidatus* Bathyarchaeota archaeon isolate PFL13 seq_4456982. The protein ID is MCJ7457314.1. In the MAG of a *Candidatus* Bathyarchaeota archaeon, a tandem array of the gene encoding ribulose-1,5-bisphosphate isomerase (abbreviated as R15P1; colored in green), which is involved in the PBP pathway, is found adjacent to the form Iα gene. Dotted lines with arrowheads at both edges correspond to the contig ends. Abbreviations: PRK, phosphoribulokinase; FBP, fructose-1,6-bisphosphate aldolase/phosphatase; RuPE, ribulose-phosphate 3-epimerase; RPI, ribose-5-phosphate isomerase; TKT, transketolase; FBA, fructose-bisphosphate aldolase; GAPDH, glyceraldehyde-3-phosphate dehydrogenase; PGK, phosphoglycerate kinase.

No experiment is currently available to examine whether the archaeal form Iα RuBisCO is involved in the PBP pathway in the particular archaeon. To examine the putative function of the archaeal form Iα RuBisCO, we first need to establish its laboratory culture, or those of archaea whose genomes harbor the operon encoding form Iα RuBisCO and R15P isomerase, if it exists.

### Evolutionary trajectory of the archaeal form Iα RuBisCO

Metagenomic data have been rapidly accumulated in public databases and are undoubtedly significant in understanding the diversity of organisms that thrive in nature. Intriguingly, no study had reported any form I-related RuBisCO in Archaea prior to this study. Depending on how frequently future studies retrieve form I-related RuBisCO sequences from archaeal MAGs, two scenarios are possible for the evolutionary trajectories of the archaeal form Iα RuBisCO gene identified in this study. The first scenario posits a primary absence of form I-related RuBisCO in Archaea, and thus, this type of RuBisCO identified in archaeal genomes (including the form Iα identified in this study) resulted from the horizontal transfer of bacterial genes. We currently favor a horizontal origin for the particular form Iα gene in the archaeal MAG, and additional form I-related RuBisCO genes may be found in as-yet-unstudied archaeal genomes. To strengthen this scenario, it would be ideal to pinpoint the bacterial donor of the RuBisCO gene. As this issue is left unresolved in the current RuBisCO phylogeny, ongoing sequencing of bacterial MAGs is necessary to improve the diversity of form I-related RuBisCO in Bacteria.

We cannot definitively rule out the possibility that this type of RuBisCO is native to at least a subtree of Archaea. The second scenario will be favored if form I-related RuBisCO is identified in a wide range of Archaea lineages in future metagenomic analyses. In summary, to evaluate the two scenarios for form I-related RuBisCO in Archaea, we require a set of this type of RuBisCO that encompasses more phylogenetically diverse species in Bacteria and Archaea than those analyzed in this study.

## Conclusion

In this study, we evaluated issues in the evolution of form I-related RuBisCOs based on their updated diversity. The detailed phylogenetic analyses conducted here demonstrated that neither monophyly of the form I′ sequences nor form Iα sequences is conclusive. These issues are left for future studies that incorporate a wider collection of form I-related sequences than those considered in the analyses presented here. Most notably, we identified the first form Iα RuBisCO gene in an archaeal MAG, and the gene product likely participates in the PBP pathway in the corresponding archaeon. In sum, the phylogenetic distribution of each RuBisCO form related to form I appeared to be less strict than we previously thought. It is also less obvious which pathway a particular RuBisCO form is involved in, as a form III RuBisCO was reported to be involved in the CBB pathway (Frolov et al. 2019), and a form Iα RuBisCO likely occupies a part of the PBP pathway (this study).

## Acknowledgements

Computations were partially performed on the NIG supercomputer at ROIS National Institute of Genetics, Mishima, Japan. This work was supported by grants from the Japanese Society for Promotion of Sciences (No. 23H02535 awarded to Y.I. and 24K09587 awarded to T.N.).

## Data availability

The RuBisCO alignments analyzed in this study, and the Newick-formatted treefiles for the ML trees inferred from these alignments are available upon request to the corresponding author.

## Notes

https://drive.google.com/drive/folders/1jNvr1gsqDeEgZczHAsGQzZ5tOZbA98rb?usp=sharing

### Competing Interest Statement

The authors have declared no competing interest.

## References

Andersson, I. and Taylor, T.C. (2003) Structural framework for catalysis and regulation in ribulose-1,5-bisphosphate carboxylase/oxygenase. Arch Biochem Biophys 414:130–140.

Aono, R., Sato, T., Yano, A., Yoshida, S., Nishitani, Y., Miki, K., Imanaka, T., and Atomi, H. (2012) Enzymatic characterization of AMP phosphorylase and ribose-1,5-bisphosphate isomerase functioning in an archaeal AMP metabolic pathway. J Bacteriol 194:6847–6855.

Aono, R., Sato, T., Imanaka, T., and Atomi, H. (2015) A pentose bisphosphate pathway for nucleoside degradation in archaea. Nat Chem Biol 11:355–360.

Ashida, H., Saito, Y., Kojima, C., Kobayashi, K., Ogasawara, N., and Yokota, A. (2003) A functional link between RuBisCO-like protein of Bacillus and photosynthetic RuBisCO. Science 302:286–290.

Bamba, K., Harada, R., and Inagaki, Y. (2024) AUTOEB: A software for systematically evaluating bipartitions in a phylogenetic tree employing an approximately unbiased test. IPSJ Trans Bioinformat 17:72–82.

Banda, D.M., Pereira, J.H., Liu, A.K., Orr, D.J., Hammel, M., He, C., Parry, M.A.J., Carmo-Silva, E., Adams, P.D., Banfield, J.F., and Shih, P.M. (2020) Novel bacterial clade reveals origin of form I RuBisCO. Nat Plants 6:1158–1166.

Bar-On, Y.M. and Milo, R. (2019) The global mass and average rate of rubisco. Proc Natl Acad Sci USA 116:4738–4743.

Camacho, C., Coulouris, G., Avagyan, V., Ma, N., Papadopoulos, J., Bealer, K., and Madden, T.L. (2009) BLAST+: architecture and applications. BMC Bioinformat 10:421.

Capella-Gutiérrez, S., Silla-Martínez, J.M., and Gabaldón, T. (2009) trimAl: a tool for automated alignment trimming in large-scale phylogenetic analyses. Bioinformatics 25:1972–1973.

Criscuolo, A. and Gribaldo, S. (2010) BMGE (Block Mapping and Gathering with Entropy): a new software for selection of phylogenetic informative regions from multiple sequence alignments. BMC Evol Biol 10:210.

Erb, T.J., Evans, B.S., Cho, K., Warlick, B.P, Sriram, J., Wood, B.M., Imker, H.J., Sweedler, J.V., Tabita, F.R., and Gerlt, J.A. (2012) A RuBisCO-like protein links SAM metabolism with isoprenoid biosynthesis. Nat Chem Biol 8:926–32.

Erb, T.J. and Zarzycki, J. (2018) A short history of rubisco: the rise and fall (?) of Nature’s predominant CO2 fixing enzyme. Curr Opin Biotechnol 49:100–107.

Ezaki, S., Maeda, N., Kishimoto, T., Atomi, H., and Imanaka, T. (1999) Presence of a structurally novel type ribulose-bisphosphate carboxylase/oxygenase in the hyperthermophilic archaeon, Pyrococcus kodakaraensis KOD1. J Biol Chem 274:5078–5082.

Felsenstein, J. (1985) Confidence limits on phylogenies: An approach using the bootstrap. Evolution 39:783–791.

Frolov, E.N., Kublanov, I.V., Toshchakov, S.V., Lunev, E.A., Pimenov, N.V., Bonch-Osmolovskaya, E.A., Lebedinsky, A.V., and Chernyh, N.A. (2019) Form III RuBisCO-mediated transaldolase variant of the Calvin cycle in a chemolithoautotrophic bacterium. Proc Natl Acad Sci USA 116:18638–18646.

Guindon, S., Dufayard, J.F., Lefort, V., Anisimova, M., Hordijk, W., and Gascuel, O. (2010) New algorithms and methods to estimate maximum-likelihood phylogenies: assessing the performance of PhyML 3.0. Syst Biol 59:307–321.

Hanson, T.E., and Tabita, F.R. (2003) Insights into the stress response and sulfur metabolism revealed by proteome analysis of a Chlorobium tepidum mutant lacking the Rubisco-like protein. Photosynth Res 78:231–248.

Heath, T., Medtke, S.M., and Hillis, D.M. (2008) Taxon sampling and the accuracy of phylogenetic analyses. J Systmat Evol 46:239–257.

Hoang, D.T., Chernomor, O., von Haeseler, A., Minh, B.Q., and Vinh, L.S. (2018) UFBoot2: Improving the ultrafast bootstrap approximation. Mol Biol Evol 35:518–522.

Huang, Y., Niu, B., Gao, Y., Fu, L., and Li, W. (2010) CD-HIT Suite: a web server for clustering and comparing biological sequences. Bioinformatics 26:680–682.

Huang, Q. and Szebenyi, D.M.E. (2023) Crystal structure of a type III Rubisco in complex with its product 3-phosphoglycerate. Proteins 91:330–337.

Isogai, R., Harada, R., Nakayama, T., and Inagaki, Y. (2025) Phylogenetic dissection provides insights into the incongruity in the tree of Archaeplastida between the analyses of nucleus- and plastid-encoded proteins. bioRχiv 10.1101/2025.05.06.652364.

Kalyaanamoorthy, S., Minh, B.Q., Wong, T.K.F., von Haeseler, A., and Jermiin, L.S. (2017) ModelFinder: fast model selection for accurate phylogenetic estimates. Nat Methods 14:587–589.

Katoh, K. and Standley, D.M. (2013) MAFFT multiple sequence alignment software version 7: improvements in performance and usability. Mol Biol Evol 30:772–80.

Kishino, H. and Hasegawa, M. (1989) Evaluation of the maximum likelihood estimate of the evolutionary tree topologies from DNA sequence data, and the branching order in Hominoidea. J Mol Evol 29:170–179.

Kitano, K., Maeda, N., Fukui, T., Atomi, H., Imanaka, T., and Miki, K. 2001. Crystal structure of a novel-type archaeal Rubisco with pentagonal symmetry. Structure 9:473–481.

Liu, A.K., Kaeser, B., Chen, L., West-Roberts, J., Taylor-Kearney, L.J., Lavy, A., Günzing, D., Li, W.J., Hammel, M., Nogales, E., Banfield, J.F., and Shih, P.M. (2023) Deep-branching evolutionary intermediates reveal structural origins of form I RuBisCO. Curr Biol 33:5316–5325.e3.

Maeda, N., Kitano, K., Fukui, T., Ezaki, S., Atomi, H., Miki, K., and Imanaka, T. (1999) Ribulose bisphosphate carboxylase/oxygenase from the hyperthermophilic archaeon Pyrococcus kodakaraensis KOD1 is composed solely of large subunits and forms a pentagonal structure. J Mol Biol 293:57–66.

Minh, B.Q., Nguyen, M.A.T., and von Haeseler, A. (2013) Ultrafast approximation for phylogenetic bootstrap. Mol Biol Evol 30:1188–1195.

Minh, B.Q., Schmidt, H.A., Chernomor, O., Schrempf, D., Woodhams, M.D., von Haeseler, A., and Lanfear, R. (2020) IQ-TREE 2: New models and efficient methods for phylogenetic inference in the genomic era. Mol Biol Evol 37:1530–1534.

Nabhan, A.R. and Sarkar, I.N. (2011) The impact of taxon sampling on phylogenetic inference: A review of two decades of controversy. Brief Bioinformat 13:122–134.

Prywes, N., Phillips, N.R., Tuck, O.T., Valentin-Alvarado, L.E., and Savage, D.F. (2023) Rubisco function, evolution, and engineering. Ann Rev Biochem 92:385–410.

Satagopan, S., Chan, S., Perry, L.J., and Tabita, F.R. (2014) Structure-function studies with the unique hexameric form II ribulose-1,5-bisphosphate carboxylase/oxygenase (Rubisco) from Rhodopseudomonas palustris. J Biol Chem 289:21433–21450. 10.1074/jbc.M114.578625.

Sato, T., Atomi, H., and Imanaka, T. (2007) Archaeal type III RuBisCOs function in a pathway for AMP metabolism. Science 315:1003–1006.

Schneider, G., Lindqvist, Y., Brändén, C., and Lorimer, G. 1986. Three-dimensional structure of ribulose-1,5-bisphosphate carboxylase/oxygenase from Rhodospirillum rubrum at 2.9 Å resolution. EMBO J 5:3409–3415.

Schulz, L., Guo, Z., Zarzycki, J., Steinchen, W., Schuller, J.M., Heimerl, T., Prinz S., Mueller-Cajar, O., Erb, T.J., and Hochberg, G.K.A. (2022) Evolution of increased complexity and specificity at the dawn of form I Rubiscos. Science 378:155–160.

Shimodaira, H., and Hasegawa, M. (1999) Multiple comparisons of log-likelihoods with applications to phylogenetic inference. Mol Biol Evol 16:1114– 1116.

Shimodaira, H. (2002) An approximately unbiased test of phylogenetic tree selection. Syst Biol 51:492–508.

Tabita, F.R., Satagopan, S., Hanson, T.E., Kreel, N.E., and Scott, S.S. (2008) Distinct form I, II, III, and IV Rubisco proteins from the three kingdoms of life provide clues about Rubisco evolution and structure/function relationships. J Experiment Bot 59:1515–1524.

Varaljay, V.A., Satagopan, S., North, J.A., Witte, B., Dourado, M.N., Anantharaman, K., Arbing, M.A., McCann, S.H., Oremland, R.S., Banfield, J.F., Wrighton, K.C., and Tabita, F.R. (2016) Functional metagenomic selection of ribulose 1, 5-bisphosphate carboxylase/oxygenase from uncultivated bacteria. Environ Microbiol 18:1187– 1199.

Wang, H.C., Minh, B.Q., Susko, E., and Roger, A.J. (2018) Modeling site heterogeneity with posterior mean site frequency profiles accelerates accurate phylogenomic estimation. Syst Biol 67:216–235.

Watson, G.M.F., and Tabita, F.R. (1997) Microbial ribulose 1,5-bisphosphate carboxylase/oxygenase: a molecule for phylogenetic and enzymological investigation, FEMS Microbiol Lett 146:13–22.

West-Roberts, J.A., Matheus-Carnevali, P.B., Schoelmerich, M.C., Al-Shayeb, B., Thomas, A.D., Sharrar, A., He, C., Chen, L-X., Lavy, A., Keren, R., Amano, Y., and Banfield, J.F. (2021) The Chloroflexi supergroup is metabolically diverse and representatives have novel genes for non-photosynthesis based CO2 fixation. bioRχiv 10.1101/2021.08.23.457424.

Wrighton, K.C., Castelle, C.J., Varaljay, V.A., Satagopan, S., Brown, C.T., Wilkins, M.J., Thomas, B.C., Sharon, I., Williams, K.H., Tabita, F.R., and Banfield, J.F. (2016) RubisCO of a nucleoside pathway known from Archaea is found in diverse uncultivated phyla in bacteria. ISME J 10:2702– 2714.

Yang, Z., Zeng, X., Zhao, Y., Chen, R. (2023) AlphaFold2 and its applications in the fields of biology and medicine. Sig Transduct Target Ther 8:115.

Yazaki, E., Yabuki, A., Imaizumi, A., Kume, K., Hashimoto, T., Inagaki, Y. (2022) The closest lineage of Archaeplastida is revealed by phylogenomics analyses that include Microheliella maris. Open Biol 12:210376.

